# Improving SBRT by re-wiring immunosuppressive neutrophils in murine pancreatic ductal adenocarcinoma

**DOI:** 10.1101/2025.01.10.632412

**Authors:** Joseph D. Murphy, Gary Hannon, Bradley N. Mills, Angela L. Hughson, Taylor P. Uccello, Nicholas W. Gavras, Jian Ye, Brian A. Belt, Maggie L. Lesch, Sarah L. Eckl, Edith M. Lord, Minsoo Kim, Haoming Qiu, David C. Linehan, Scott A. Gerber

**Affiliations:** Department of Microbiology and Immunology, University of Rochester Medical Center, Rochester, NY, USA; Department of Surgery, University of Rochester Medical Center, Rochester, NY, USA; Department of Radiation Oncology, University of Rochester Medical Center, Rochester, NY, USA

## Abstract

Radiation is used to treat pancreatic ductal adenocarcinoma (PDAC) in the locally-advanced setting. Stereotactic body radiation therapy (SBRT), in particular, has shown improved outcomes against conventional RT in a number of clinical trials. One key cell type involved in the response to RT are neutrophils. These innate immune cells are the first responders to tissue damage and infiltrate irradiated tumors in high numbers to rectify injury. Neutrophils typically possess an immunosuppressive, wound healing phenotype in this scenario, which allows the tumor to recover and actively suppress immunological efforts to eradicate the disease. Here, we aimed to elucidate the role of neutrophils in a murine model of PDAC treated with SBRT. Mice harboring PDAC tumors were treated with targeted SBRT and neutrophils were determined to be significantly increased in the tissue when compared to unirradiated controls. Additionally, phenotypic analysis determined that these cells were largely immunosuppressive and depletion studies confirmed they played a key role in acquired radioresistance in our model. In order to establish whether these infiltrating cells could be re-wired to contribute to anti-tumor immunity, we utilized a novel combination therapy consisting of SBRT and microspheres containing recombinant IL-12 to attempt to repolarize these cells. Transcriptomic analysis confirmed intratumoral neutrophils underwent considerable changes indicative of an immunostimulatory, anti-tumor phenotype following treatment. Moreover, depleting these cells resulted in a loss of treatment efficacy, suggesting that neutrophil re-wiring was vital for the therapeutic outcome. This study highlights neutrophils as key players in SBRT and IL-12 treatment and confirms they can act as a double-edged sword depending on the treatment employed.

## 1. Introduction

Radiation is an approved treatment modality for locally-advanced pancreatic ductal adenocarcinoma (PDAC). Historically, this treatment has been delivered through low dose fractions (1.8-2 Gy) over 5-6 weeks [1]. This conventional regimen has largely been replaced by stereotactic body radiation therapy (SBRT), which enables precise delivery of higher dose oligofractions (5-8 Gy) over shorter periods (1-2 weeks) and has proven a safer and more efficacious alternative across multiple preclinical and clinical studies [2-4].

Neutrophils are typically the first line of defence for sites of infection and tissue damage [5]. In cancer, they can exhibit an immune stimulatory or immunosuppressive phenotype depending on the context of the tumor microenvironment [6]. Generally, however, higher levels of tumor-associated neutrophils (TANs) are considered a marker of poorer prognosis across multiple solid malignancies [7]. Following radiation, neutrophils are recruited to tumors in high numbers as an immunological response to tissue damage. Once there, these cells largely exhibit an immunosuppressive wound-healing phenotype that contributes to many pro-tumor responses, including acquired radioresistance [8-10]. Multiple studies have sought to repolarize these cells or mitigate their tumor infiltration to enhance the efficacy of immunotherapies and chemotherapies in PDAC [11-13]. Despite this, targeting these cells in the context of RT is underdeveloped.

Here, we aimed to decipher the role of neutrophils in SBRT treatment of murine PDAC. Neutrophil recruitment to these tumors following SBRT was confirmed by flow cytometry and immunohistochemistry (IHC) along with their immunosuppressive phenotype, and depletion studies noted their key role in treatment response. To determine whether radio-responsive neutrophils could be re-wired towards an anti-tumor phenotype, we utilized a combination treatment of SBRT and IL-12 microspheres (MS), which we recently employed to eliminate primary PDAC disease and repolarize immunosuppressive macrophage and monocyte populations [2]. This combination treatment is also currently undergoing clinical evaluation at our medical center for locally-advanced PDAC (NCT06217666). Bulk RNA sequencing of neutrophils following treatment confirmed profound transcriptomic changes suggestive of a more anti-tumor, immunostimulatory phenotype. Moreover, subsequent depletion studies also confirmed neutrophils were essential to treatment efficacy. This study highlights neutrophils as major players governing radiation responses in PDAC. Re-wiring this immunosuppressive population can sensitize this cancer to a variety of therapeutic modalities, including SBRT.

## 2. Methods

### 2.1 Cell culture

KCKO cells (derived from P48-Cre/LSL-Kras^G12D^ C57BL/6 mice) were maintained in MAT/P (US Patent number 4.816.401) supplemented with 5% fetal bovine serum (FBS; GIBCO), and 1% penicillin/streptomycin (Thermo Fisher Scientific). This cell line was provided by Dr. David DeNardo (Washington University, St. Louis) and were stably transfected to express firefly luciferase (KCKO-luc).

### 2.2 *In vivo* animal studies

All experiments were approved by the University Committee on Animal Resources (UCAR) and were performed in compliance with both the National Institute of Health (NIH) and University approved guidelines for the care and use of animals. 6-8-week-old female C57BL/6J mice (Jackson Laboratory) were used for this study. All mice were subjected to a twelve-hour light/dark cycle and kept in individually ventilated cages with bedding and nesting material.

### 2.3 Orthotopic tumor mouse model

6-8-week-old female C57BL/6 mice were anesthetized with vaporized isoflurane (VetFlo, Kent Scientific) and a 10-mm laparotomy incision was made to expose the pancreas. 100,000 KCKO-luc cells prepared in 40 μl mixture of 1:1 PBS:Matrigel (BD Bioscience) were injected into the tail of the pancreas. Next, two 4-mm titanium fiducial clips (Weck) were placed on both flanks of the Matrigel bubble to facilitate radiation targeting. The peritoneal cavity was then sutured using 4-0 Vicryl SH-1 (eSutures) and the skin was stapled using 9-mm wound clips (Reflex).

### 2.4 *In vivo* radiation treatment

SBRT was delivered using the Small Animal Radiation Research Platform (SARRP, XStrahl) using a 5 mm collimator. Mice were anaesthetized with vaporized isoflurane before and during radiation treatment. Tumor specific delivery was achieved using titanium fiducial markers visualized through a pre-treatment computed tomography (CT) scan. SBRT was administered following a schedule of 6 Gy radiation in 4 fractions on days 6-9 post-implantation.

### 2.5 Bioluminescent tracking of tumor growth

Mice bearing KCKO-luc tumors were anesthetized with vaporized isoflurane and injected subcutaneously with D-luciferin (2.5mg, Invitrogen) in 100 μL PBS. Mice were then placed in the IVIS Spectrum Imaging System (IVIS, PerkinElmer) in the right lateral recumbent position on a platform heated to 37° C. Twelve consecutive images were taken at 2-minute intervals for 24 minutes. Bioluminescence (BLI; p/sec/cm^2^/sr) was calculated using standardized regions of interest (ROIs) manually placed over peak bioluminescent signals. Peak intensity was recorded for each tumor.

### 2.6 Flow cytometry

Tumor-bearing mice were sacrificed, and tumors were harvested, weighed, and processed using a scalpel in 30% collagenase in PBS with 1:1000 brefeldin added (Sigma Aldrich). The Macs homogenizer (Miltenyl Biotec) was used three times using the tumor setting with 10 minute incubations in a water bath at 37° C between every run. Tumors were washed twice with PBS containing 0.1% sodium azide, 1% bovine serum albumin (PAB) and brefeldin (at 1:1000 dilution) and filtered through a 70 μm filter. 1×10^6^ cells per sample were stained for surface and intracellular markers using a range of antibodies including CD45-PerCPcy5.5 (30-f11, BD), CD11b-BV510 (M1/70, BD), Ly6C-PEcy7 (HK1.4, Biolegend), Ly6G-BV605 (1A8, BD), I-A/I-E-APC (M5/114.15.2, Biolegend), F4/80-APC (BM8, Biolegend), CD8-PEcy5 (53-6.7, BD), CD4-BUV395 (RM4-5, BD), CD19-BV786 (ID3, BD), Siglec-F-AF488 (1RNM44N, eBioscience), Arg-1-AF700 (A1exF5, eBioscience), SPARC-AF405 (124413, R&D) and IFNγ-BV650 (XMG1.2, BD). Samples were stained for 60 minutes at 4 °C for surface markers, then washed and resuspended in CytoPerm/CytoFix (BD Bioscience) for twenty minutes at 4 °C for downstream intracellular staining protocols. For intracellular staining, IFNγ BD Cytofix/Cytoperm Plus kit (BD) was used as per manufacturer protocol. 100,000 events were collected using BD LSRII, BD FACS Symphony A1 or BD LSR Fortessa.

### 2.7 Immunohistochemistry

Tumors were harvested and placed in 10% neutral buffered formalin (Azer Scientific). Tumor samples were formalin-fixed, paraffin-embedded and sectioned using a microtome to generate 5-μm sections placed on glass slides. A sample from each tumor was stained with Mayer’s hematoxylin and Eosin (H&E; Richard Allan Scientific) by the Wilmot Cancer Institute Histopathology core facility. For immunohistochemistry (IHC), samples were deparaffinized, rehydrated and baked with sodium citrate buffer for antigen retrieval, before being stained with a rabbit anti-mouse Ly6G antibody (1A8, Abcam) followed by a polymer based anti-Rabbit detection kit (Abcam) and counter stained with hematoxylin. Stained slides were scanned using the Aperio Slide Scanner (Leica). Strong positive pixel number was calculated using Aperio Image Scope software (Leica).

### 2.8 Luminex protein quantification assay

Mouse tumors were harvested and dissociated with a tissue homogenizer in 5 ml of 0.5x Lysis Buffer II (R&D Systems) with added 1x Halt Protease Inhibitor and 1x Phosphatase Inhibitor Cocktail (Thermo Fisher). Tissues were lysed at room temperature for 30 min on a plate shaker at 80 rpm. Luminex Assays were performed using a Multi-Analyte Kit (R&D Systems). Pierce BCA Protein Assays (ThermoFisher Scientific) were performed to normalize data to total protein concentrations.

### 2.9 Neutrophil depletion

To deplete neutrophils, mice were injected subcutaneously with 200 μg rat anti-mouse Ly6G (1A8 clone, BioXCell) or rat IgG2a isotype control in 100 μl of PBS every three days from day 5 onwards (the first day of SBRT), and doses were continued thereafter three times per week until end of study.

### 2.10 *In vivo* microsphere treatment

One day post the final fraction of SBRT, tumor-bearing mice were anesthetized with vaporized isoflurane (VetFlo, Kent Scientific) and a 10-mm laparotomy incision was made to expose the pancreatic tumor. Empty MS control (2 mg polylactic acid beads) or IL-12 MS (2 mg polylactic acid beads containing 0.5 mg recombinant IL-12) were injected at two separate intratumoral locations with 10 μl per injection site using a 32-gauge Hamilton syringe. The peritoneal cavity was sutured and skin stapled as mentioned previously. Microspheres were created utilizing the reported phase inversion technique [14, 15].

### 2.11 RNA Sequencing

Mice were sacrificed and three mouse tumors were pooled per treatment group. Tissues were harvested, weighed, and mechanically dissociated as before. Tumors were washed twice using PBS and passed through a 70-μm filter. 5×10^6^ cells per sample were stained for various surface markers using a range of antibodies for 60 minutes at 4°C in the dark. Ly6G^hi^ neutrophils were sorted on a FACSAria II cell sorter (BD Biosciences) using a 100 μm nozzle. Cells were lysed in Buffer RLT containing β-mercaptoethanol and homogenized via QIAShredder spin columns. RNA was purified using the RNeasy Micro Kit (QIAGEN). Downstream RNA sequencing and analysis was performed by the University of Rochester Genomics Research Center. RNA quality was assessed using an Agilent Bioanalyzer (Agilent), with all samples demonstrating RNA integrity values > 5. cDNA libraries were constructed using the TruSeq RNA Sample Preparation Kit V2 (Illumina) following manufacturer’s instructions, and sequencing was performed using an Illumina high-throughput HiSeqTM 2500 platform (Illumina). Differentially expressed genes (DEGs) between treatment groups compared to vehicle treated unirradiated controls were analyzed using Ingenuity Pathway Analysis (IPA) software (QIAGEN).

### 2.12 Statistical Analyses

GraphPad Prism 9 software was used for the generation of all graphs and statistical analyses. p values < 0.05 were deemed statistically significant. Tumor growth data was analyzed using Mann-Whitney U tests at each comparable timepoint or by one-way ANOVA followed by Dunnett’s test. Survival data was graphed with Kaplan-Meier curves and analyzed by the Mantel-Cox test. Statistical analysis for flow cytometry data was performed using Kruskal-Wallis or unpaired T-tests.

## 3. Results

### 3.1 Neutrophil levels increase in KCKO tumors following SBRT and correspond with temporary growth delay

We developed a clinical relevant murine model that recapitulates SBRT treatment of locally advanced PDAC. Following SBRT treatment (protocol outlined in **Figure 1A**), we determined a growth delay in the KCKO tumors that lasted until day 20 by IVIS, at which point the treated tumors returned to a growth rate matching that of unirradiated tumors (**Figure 1B**). This result is also mirrored by tumor weights harvested on days 10, 15 and 20, where a significant reduction in weight is noted at the later timepoints (**Figure 1C**). Notably, flow cytometry determined this transient growth delay corresponded with increases in tumor levels of myeloid cells (**Figure 1D**), however, macrophage levels were unchanged (**Figure 1E**), while neutrophils exhibited a delayed increased (**Figure 1F**). Interestingly, there were also no significant differences in monocytes following treatment (**Figure 1G**), suggesting neutrophils were the predominant myeloid subset that trafficked to the tumor post treatment. IHC corroborated these findings, demonstrating an increase in Ly6G+ population on day 15 (**Figure H-J**). Neutrophils in circulation express the chemokine receptor CXCR2 which binds to many chemokine ligands including CXCL1, 2 and 5. Notably, murine models of PDAC have been shown to recruit neutrophils in a CXCR2-dependent manner [16]. Luminex data showed this neutrophil infiltration corresponded with intratumoral increases in the chemokines CXCL2 and CXCL1 on days 10 and 15 respectively (**Supplementary Figure 1A & B**). Notably, no changes were observed in lymphoid populations across each timepoint (**Supplementary Figure 2A-C**).

**Figure 1.**
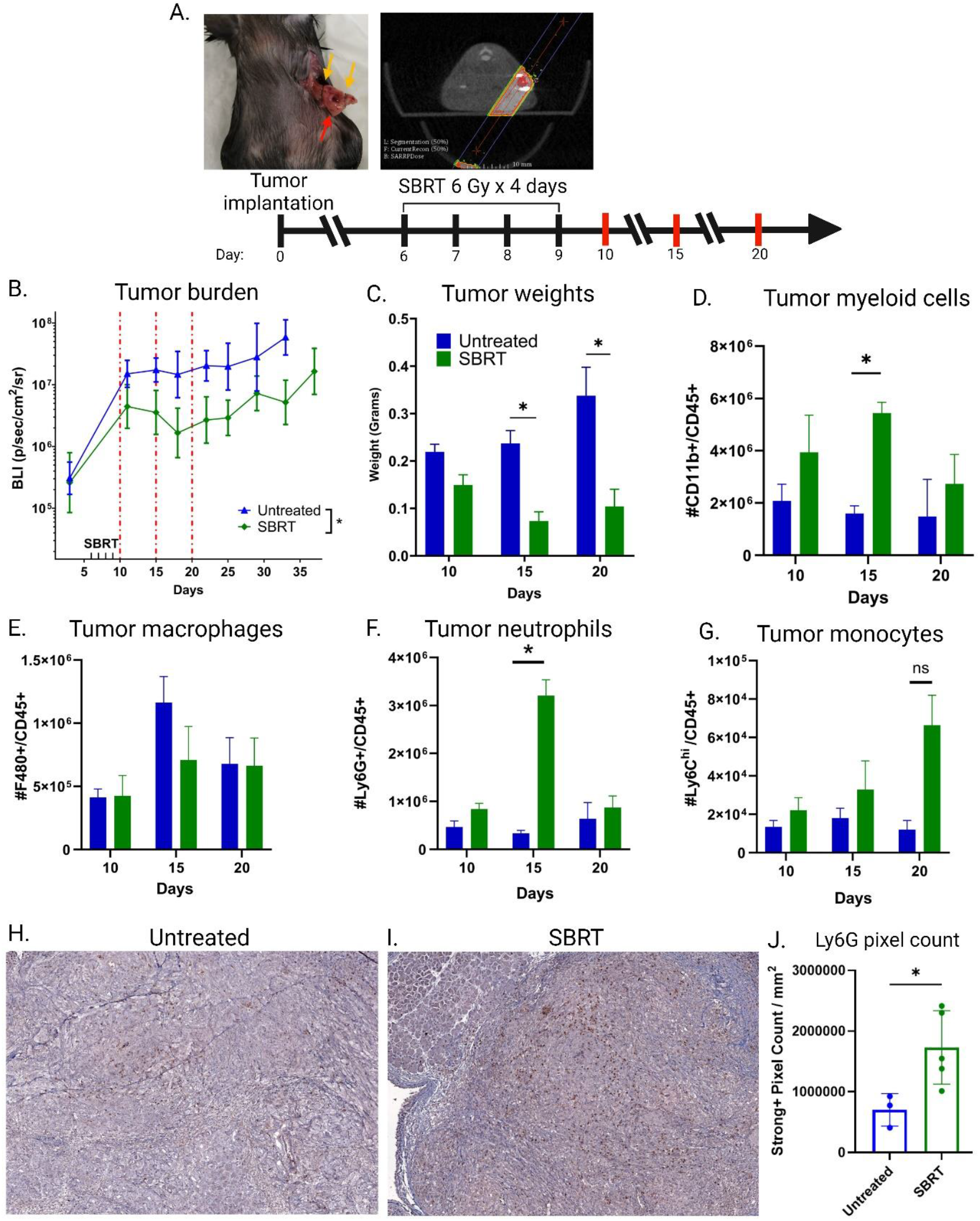
SBRT induces temporary growth delay that coincides with neutrophil infiltration into the tumor. **A**. Scheme for SBRT treatment of KCKO tumors. Red bars indicate days when tumors were harvested for weight comparison and flow cytometry analysis. Gold arrows indicate titanium fiducial clips for SBRT targeting; Red arrow indicates site of tumor injection. **B**. BLI growth curves for unirradiated and SBRT-treated animals (n=5). **C**. Tumor weights for unirradiated and SBRT-treated controls (n=5). **D-F**. Tumor levels of myeloid populations on day 10, 15 and 20. Myeloid cells (CD45+CD11b+; D), macrophages (CD45+CD11b+F4/80+; E), neutrophils (CD45+CD11b+Ly6G+; F) and monocytes (CD45+CD11b+Ly6C^HI^; G) were assessed over time by flow cytometry (n=5). **H** and **I**. Representative IHC images of Ly6G+ cells in unirradiated and SBRT-treated controls. **J**. Quantification of pixel count for Ly6G+ cells in unirradiated (n=3) and SBRT-treated (n=5) animals on day 15. *P<0.05. Data in B-G and J analyzed by Mann-Whitney U test. Data shown are mean+ SD.

### 3.2 Circulating levels of neutrophils and other myeloid cells increase following SBRT

To elucidate further the systemic response of neutrophils to radiation, we measured the density of these cells in the bone marrow, spleen and peripheral blood at the above-mentioned timepoints. Splenic myeloid cells, namely neutrophils and monocytes, increased significantly at day 10 (**Figure 2A-C**). Mirroring these findings, peripheral blood neutrophils and monocytes along with eosinophils (Siglec-F^+^) also experienced a significant rise at this timepoint (**Figure 2D-F**). Bone marrow, by contrast, displayed no changes to any of these cell types (**Figure 2H-J**).

**Figure 2.**
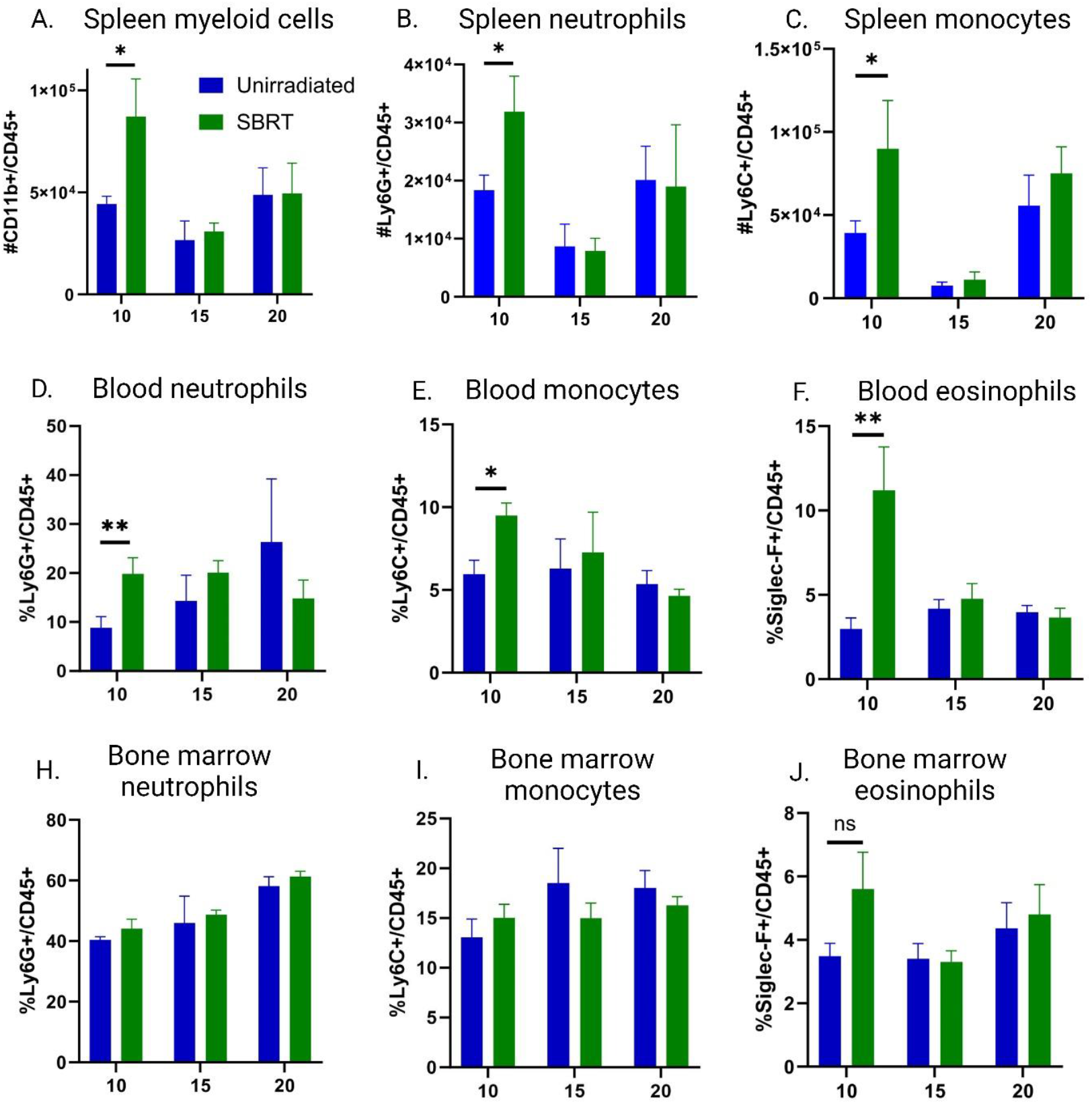
SBRT induces acute changes in myeloid levels in the blood and spleen. **A**. Splenic levels of myeloid (CD45+CD11b+), monocyte (CD45+CD11b+Ly6C^HI^) and neutrophils (CD45+CD11b+Ly6G+) over time in irradiated and SBRT-treated mice. **B**. Blood levels of monocytes, eosinophils (CD45+CD11b+Siglec-F+) over time in irradiated and SBRT-treated mice. **C**. Bone marrow levels of monocytes, eosinophils and neutrophils over time in irradiated and SBRT-treated mice (n=5). *p<0.05;**p<0.01. All data analyzed by Mann-Whitney U test. Data shown are mean+ SD.

### 3.3 Neutrophils express suppressive markers in tumors after SBRT

Given that neutrophils were upregulated in SBRT-treated tumors, we phenotyped these cells post SBRT to determine their immuno-stimulatory/suppressive phenotype following infiltration. For this, we measured MHC class II and IFN-γ expression as these are known to be markers of an anti-tumor phenotype [17]. By contrast, levels of Arg-1 and SPARC were utilized as markers of a pro-tumor phenotype [18-20]. Both unirradiated and SBRT-treated neutrophils expressed similar levels of MHC class II, whereas IFN-γ percentages were significantly increased in SBRT-treated tumors at day 15 (**Figure 3A & B**). Although IFN-γ is considered an anti-tumor cytokine, these SBRT-induced neutrophils also had significantly higher levels of Arg-1 and SPARC suggesting an overall immunosuppressive phenotype in treated tumors (**Figure 3C & D**). Taken together, these data suggest that neutrophils in SBRT treated tumors at day 15 exhibited a more suppressive phenotype with acquisition of Arg-1 and SPARC expression despite maintained expression of the pro-inflammatory IFN-γ.

**Figure 3.**
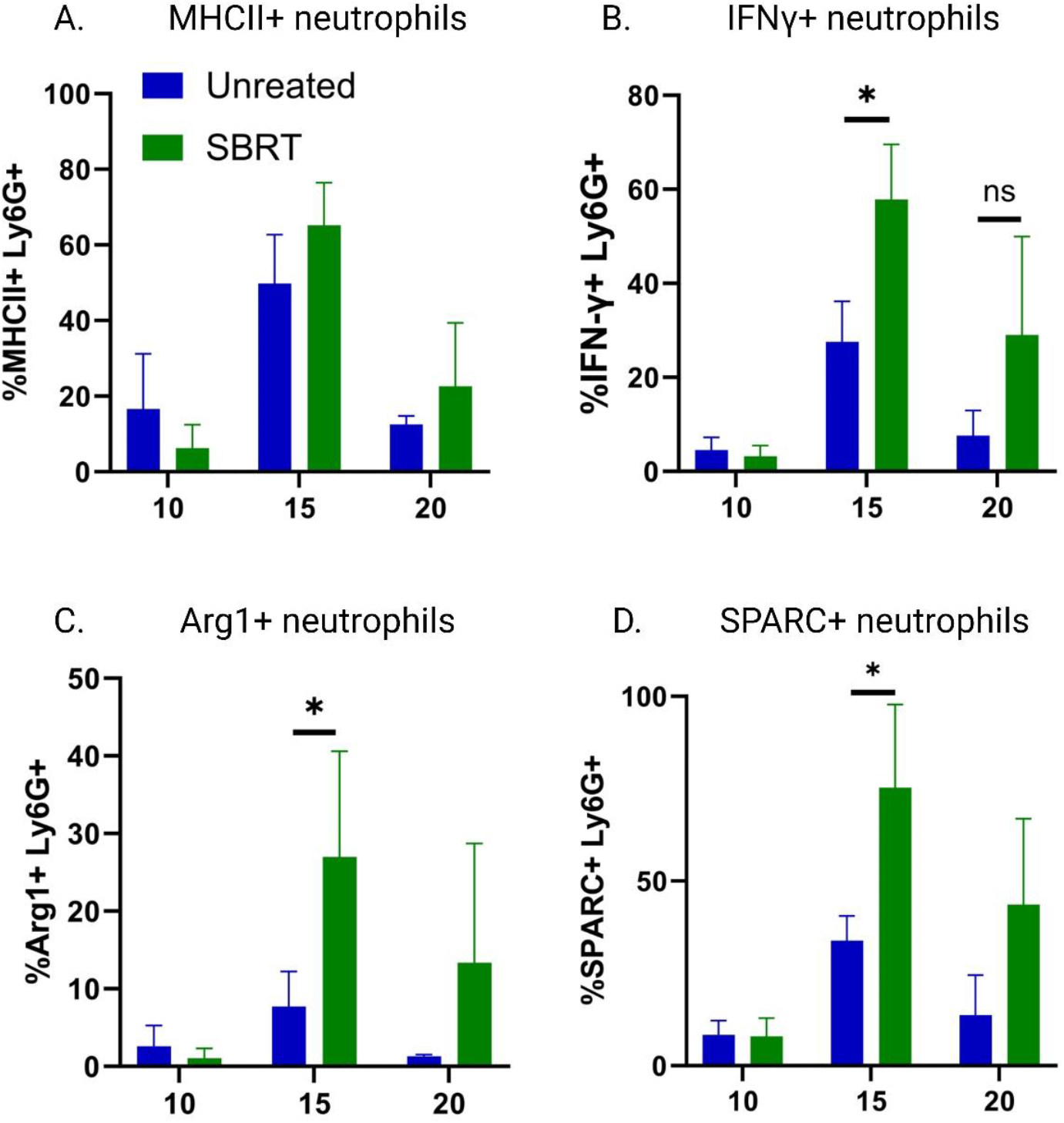
Expression of immunostimulatory and immunosuppressive markers by TANs. **A**. Expression of MHCII, **B**. IFN-γ, **C**. Arg-1 and **D**. SPARC on TANs on day 10, 15 and 20 (n=5). *p<0.05. All data analyzed by Mann-Whitney U test. Data shown are mean+ SD.

### 3.4 Depleting neutrophils during and post SBRT improves tumor control and survival

As neutrophils are increased in tumors following SBRT and display an apparent immunosuppressive phenotype, we sought to deplete these cells (via anti-Ly6G; 1A8 clone, BioXCell) to determine whether this will enhance the response to radiation (protocol outlined in **Figure 4A**). Administration of anti-Ly6G resulted in a significant reduction of intratumoral neutrophil density, and circulating levels exhibited non-significant reductions (**Figure 4B & C**). Depleting these cells during and after SBRT delayed tumor outgrowth at day 20 and extended median overall survival from 39 days in the SBRT-only group to 58 days in the depleted group (**Figure 4D & E**). These data indicate that radiation-responsive neutrophils play a role in tumor resistance to SBRT and can be targeted to enhance radiosensitivity.

**Figure 4.**
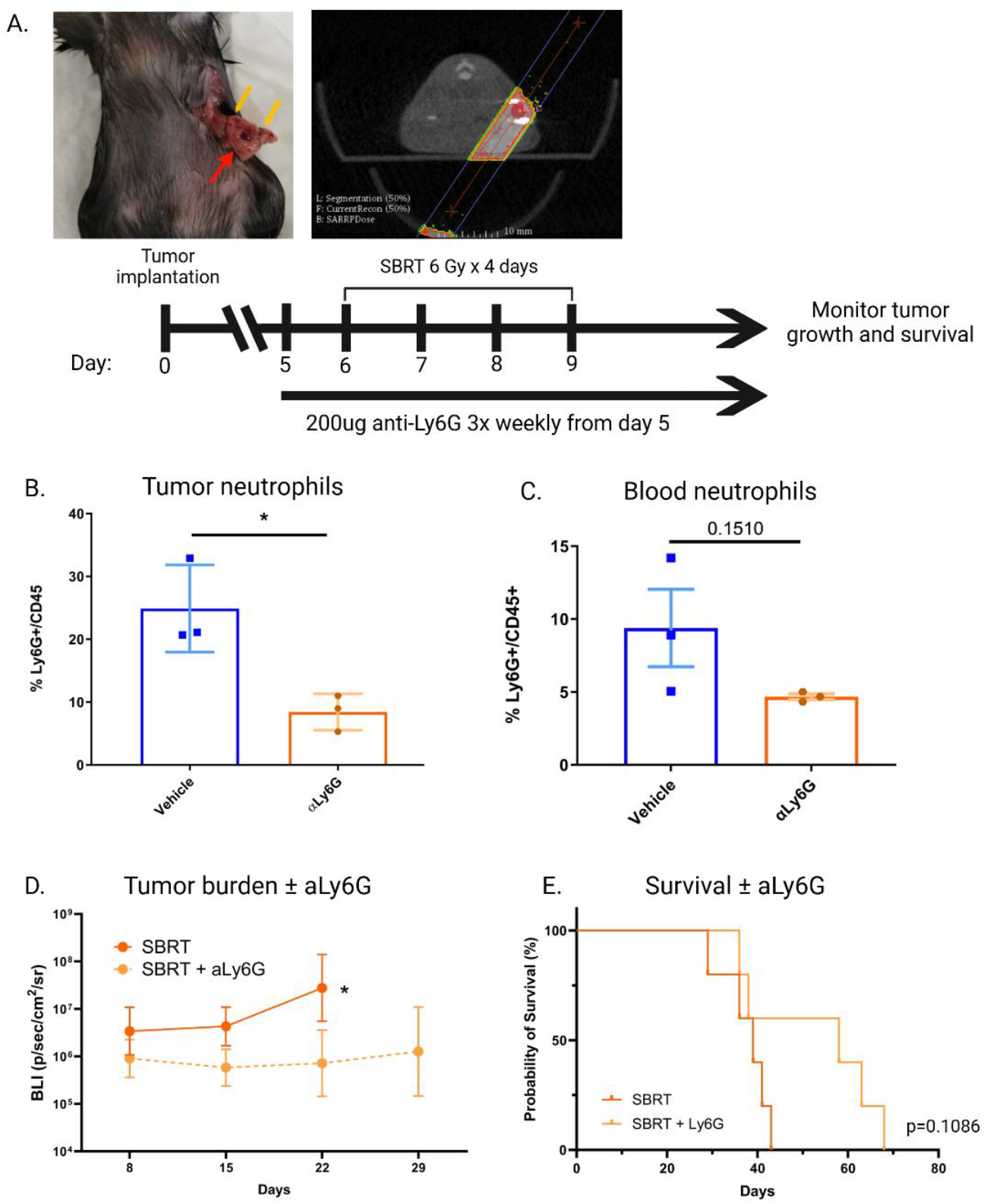
Depleting TANs can enhance the efficacy of SBRT. **A**. Scheme for SBRT and depletion treatments. **B** and **C**. Neutrophil density/concentration in the tumor and blood on day 15 in vehicle and anti-Ly6G treated animals (n=3). **D** and **E**. Growth curve and survival in vehicle and anti-Ly6G treated animals (n=5). *p<0.05. Data analyzed by Mann-Whitney U (B-D) and Mantel-Cox tests (E). Data shown are mean+ SD.

### 3.5 Re-wiring immunosuppressive neutrophils with IL-12 MS can lead to cures when combined with SBRT

Since depleting immunosuppressive neutrophils can enhance SBRT efficacy, we hypothesized that repolarizing these cells to a more anti-tumor phenotype may augment the effects of SBRT further. To achieve this, we employed a recombinant IL-12 MS treatment strategy reported by our group previously (illustrated in **Figure 5A**) [2]. Notably, delivering this cytokine via intratumoral injection one day post the last fraction of SBRT could reprogram macrophage and monocyte populations to a more pro-tumor phenotype, suggesting it could achieve similar outcomes in neutrophil populations also. As expected, and as reported previously, the combination of SBRT and IL-12 MS resulted in complete tumor rejection and 100% survival in this model (**Figure 5B & C**) [2, 21]. Additionally, bulk RNA sequencing of TANs showed a significant increase in genes associated with the IFN-γ pathway in the SBRT and IL-12 MS treated group versus unirradiated and empty MS treated controls. More specifically, a significant upregulation of genes associated with IFN-γ-induced and activated proteins such as *IFIT3, IFIT2, IFIT213, IFIT208* were observed, along with guanylate binding proteins *GBP8, GBP4, GBP6, GBP2* and XIAP Associated Factor 1 (*XAF1*) [22-25]. Likewise, these neutrophils also experienced a significant downregulation in genes associated with an immunosuppressive phenotype, such as wound-healing pathways and extracellular matrix production (*SPARC, SERPINH1, SERPINB1A, COL1A2*, and *COL3A1*) (**Figure 5D**) [20, 26-28]. Pathway analysis on the differentially expressed genes (DEG’s) between the SBRT and IL-12 MS group versus unirradiated and empty MS group showed a significant increase in pathways associated with inflammatory signalling. In particular, pathways for IFN signalling, T cell exhaustion, death receptor signalling, HMGB1 signalling, Th1 pathway and activation of pattern recognition receptors were all increased (**Figure 5E**). Of note, the majority of these gene changes were exclusive to the combination group and were not found in either single treatment group. Similarly, a volcano plot displaying log 2-fold changes in gene expressions reinforced these treatment-induced distinctions (**Figure 5F**), and a dataset for canonical myeloid-derived suppressor genes was screened to confirm that this suppressive phenotype was lost following treatment (**Figure 5G**). Hence, in response to SBRT and IL-12 treatment, neutrophils alter their transcriptome indicative of a more anti-tumor phenotype.

**Figure 5.**
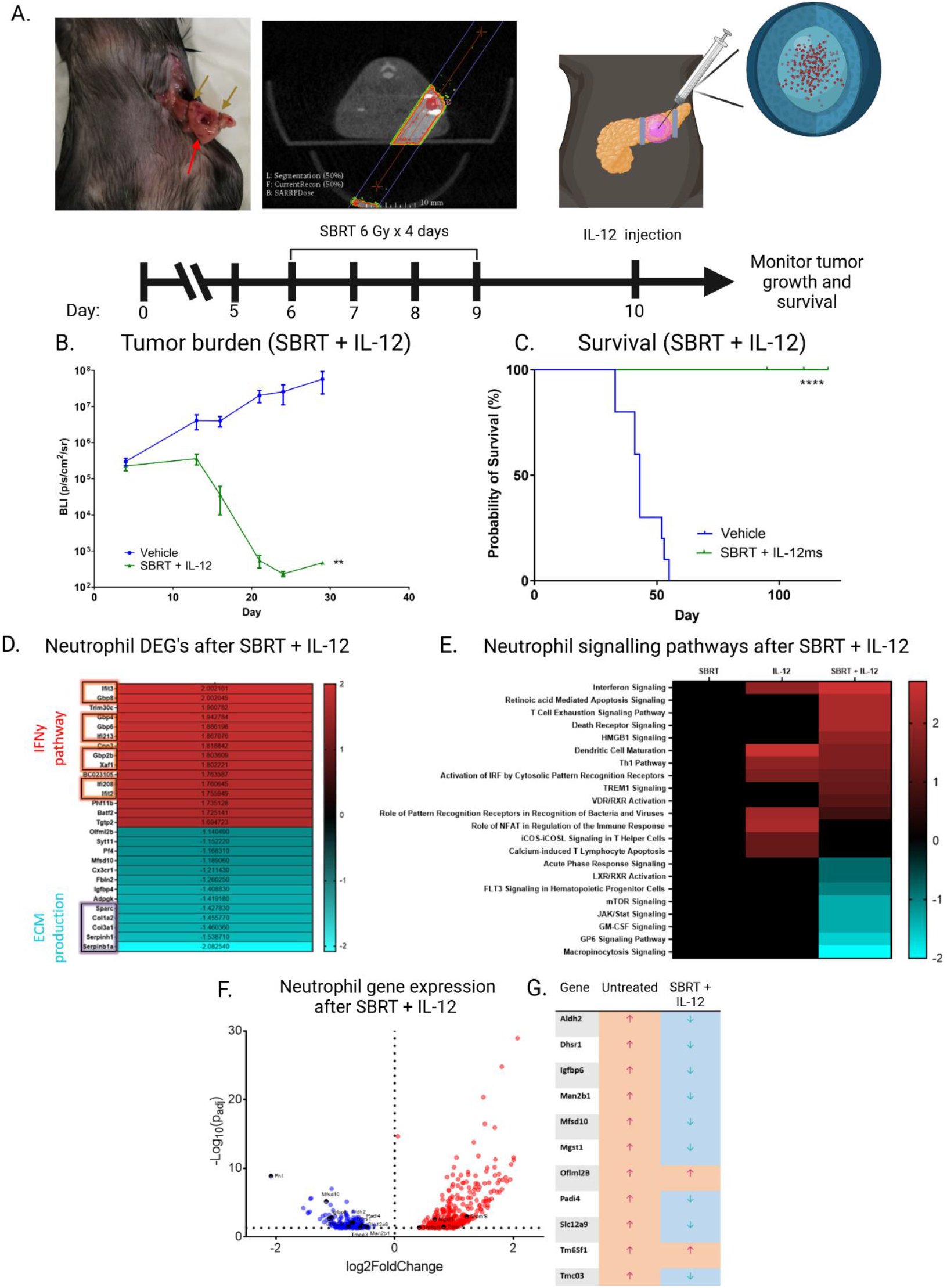
The combination of SBRT + IL-12 MS promotes pathways associated with anti-tumor repolarization. **A**. Scheme for SBRT and IL-12 MS treatment. **B**. Growth curve for SBRT and IL-12-treated and empty MS-treated, unirradiated animals (n=5-10). **C**. Survival curve for these same groups. **D**. Heat map of differentially expressed genes (DEG’s) in SBRT + IL-12 MS-treated neutrophils sorted at day 11 compared to those from unirradiated, empty microsphere treated mice (n=3). Fold change indicated within the columns and by color change. **E**. Heat map illustrating differentially expressed pathways altered in each treatment group compared to the unirradiated, empty MS control (n=3). **F**. Volcano plot of upregulated and downregulated genes in SBRT + IL-12 MS treated neutrophils compared to unirradiated, empty MS controls (n=3). **G**. Canonical genes for MDSCs were compared across treatments. **P<0.01; ****P<0.0001. Data analyzed by Mann-Whitney U (B) and Mantel-Cox (C) tests.

### 3.6 Neutrophils are required for SBRT and IL-12 MS treatment efficacy

Given that SBRT and IL-12 MS treatment could shift intratumoral neutrophils to a more anti-tumor phenotype, we next tested the contribution of these cells to overall treatment efficacy. Neutrophils were depleted during and after SBRT and IL-12 MS treatment using anti-Ly6G as described previously (**Figure 6A**). Depletion of neutrophils led to a greater tumor burden and significant decrease in survival against the combination treatment alone (**Figure 6B & C**). Notably, survival dropped to 50% after more than 100 days follow-up, demonstrating their necessity to treatment efficacy. Hence, under certain circumstances such as those generated by immunotherapies, neutrophils can become repolarized to an immunostimulatory phenotype and contribute to the anti-tumor effects of SBRT.

**Figure 6.**
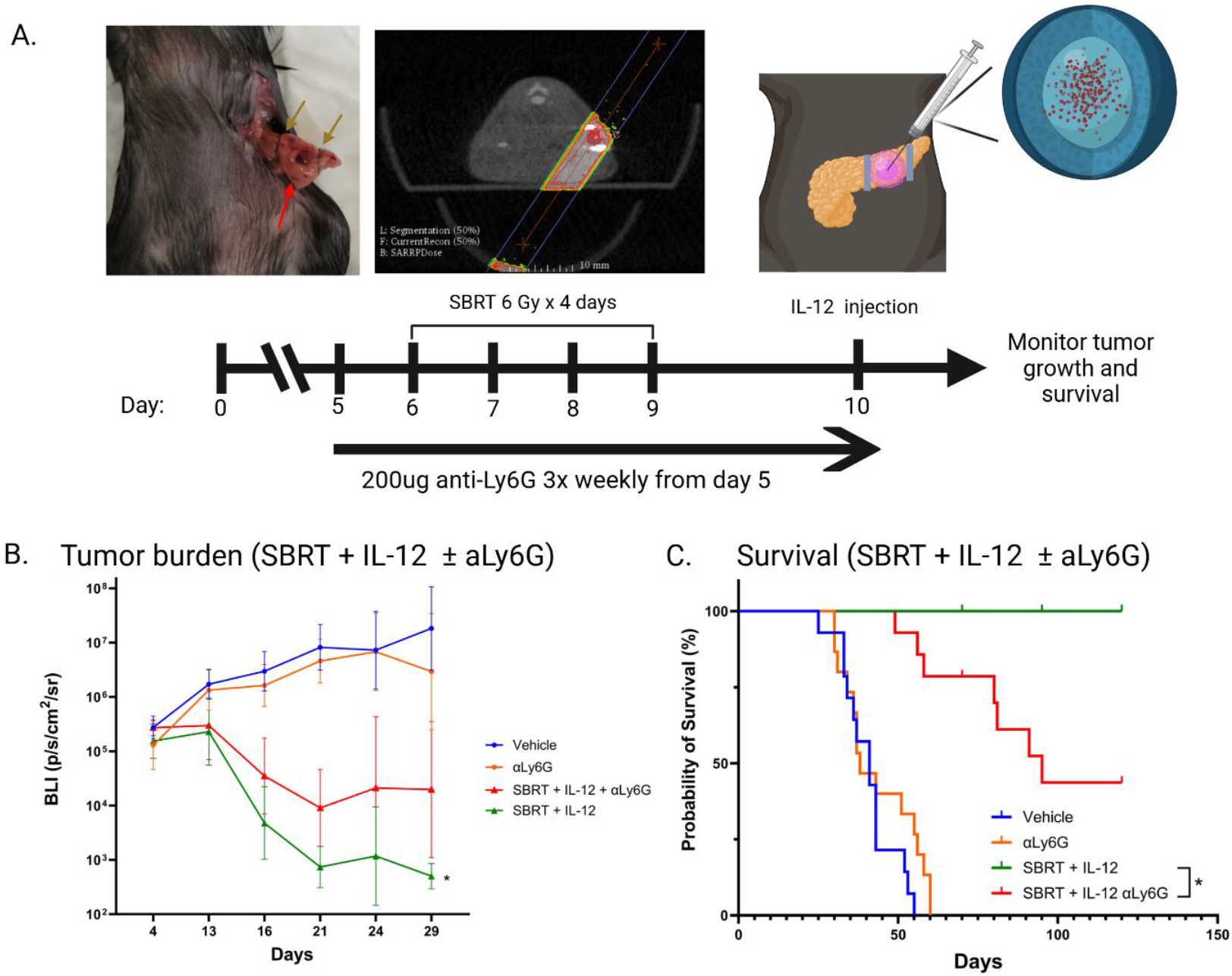
Depletion of neutrophils during SBRT + IL-12 MS treatment reduces treatment efficacy and overall survival. **A**. Treatment scheme indicating timeline of depletion along with combination SBRT + IL-12 MS treatment. **B**. BLI growth curve of each group over time. **C**. Survival curve for depleted and non-depleted groups. *p<0.05. Data analyzed by Mann-Whitney U (B) and Mantel-Cox tests (C). n=13-15. Data shown are mean+ SD.

## 4. Discussion

This study demonstrates that neutrophils are a key cell type in the response to SBRT and IL-12 MS therapy. When induced by SBRT alone, they infiltrate in high numbers and exhibit an immunosuppressive phenotype, leading to tumor outgrowth and radioresistance. Conversely, when repolarized by combination SBRT and IL-12 MS therapy they are crucial to the subsequent anti-tumor immune response. This suggests that depending on the therapeutic context, neutrophils may act as a double-edged sword, harboring a potential negative or positive prognostic outcome for patients. This is of particular significance given the numerous ongoing clinical trials involving radiation in PDAC with various combinations of immunotherapies (NCT06217666; NCT03767582; NCT05088889; NCT05721846; NCT04247165; NCT03563248). Hence, understanding the neutrophil response to these therapies may provide a framework for stratifying patients or identifying biomarkers of response in cancer.

The transient radiation response observed following SBRT involved partial tumor elimination between days 10 and 15, followed by equilibrium (shortly after day 15) and escape at day 20. This outgrowth coincided with neutrophil influx into the tumor at day 15, which we, and others, have identified as the most responsive immune cell type to SBRT [8, 29, 30]. Neutrophil recruitment to sites of infection and tissue damage is driven, in part, by chemokine signalling [31, 32]. In particular, the CXCR2 signalling axis has been shown to be essential for neutrophil recruitment to PDAC tumors [16]. Our results show that SBRT could induce intratumoral CXCL2 and CXCL1 expression at day 10 and 15 respectively, providing a chemokine gradient for neutrophil infiltration.

The role of neutrophils within irradiated tumors remains controversial as various reports describe a protumor, wound-healing, immunosuppressive neutrophil infiltrate whereas others note an antitumor, inflammatory phenotype in these cells [8, 29, 30, 33]. We identified increases in IFN-γ, SPARC and Arg-1 in irradiated tumors at day 15 in our model. While IFN-γ is a potent cytokine involved in anti-tumor immunity, it is constitutively expressed in mature neutrophils [34], and it is therefore possible that the increased number of IFN-γ^+^ neutrophils at this timepoint may be due to a retention of this cytokine and failure to release it in the TME. Nevertheless, significant increases in both SPARC and Arg-1 suggest the development of a more immunosuppressive phenotype at day 15 [20, 35]. Depleting neutrophils confirmed this immunosuppressive phenotype, as the tumor response to SBRT was more pronounced and this resulted in increases in survival.

Given these data, we sought to therapeutically reprogram these neutrophils to a more anti-tumor phenotype using polylactic acid MS containing recombinant IL-12. KCKO tumors cure by day 20 when treated with combination SBRT and IL-12 MS, in direct contrast to the tumor escape seen in SBRT alone. RNA sequencing noted large-scale changes to the transcriptome of SBRT and IL-12 MS-treated neutrophils. This is of particular interest, as neutrophils are known to have low basal levels of transcription, therefore any treatment-induced DEG’s is indicative of profound changes to this cell type. Moreover, the major pathway changes we identified were associated with the IFN response, which is crucial to the efficacy of this therapy as therapeutic effects are lost in IFN-γ-KO mice [2]. Pairing the observed increase in IFN response alongside the downregulation of genes associated with ECM remodelling, we surmise that these cells have undergone changes leading to a more anti-tumor phenotype. This also appears to be dependent on both SBRT and IL-12 treatments, as the single modalities alone had minimal (IL-12) or no effects (SBRT). Indeed, depleting these cells resulted in loss of treatment efficacy, suggesting the re-programming of these cells is a critical event governing the overall effects of this combination therapy.

## 5. Conclusion

This study demonstrates that SBRT treatment results in a transient anti-tumor effect that resolves following influx of immunosuppressive neutrophils to our PDAC murine model. Depleting these immunosuppressive neutrophils could significantly improve SBRT response, marked by greater levels of tumor control and survival. Notably, we also demonstrate that TANs could be repolarized to a more immunostimulatory phenotype following IL-12 MS treatment directly to the tumor which augments the response of SBRT, leading to cures in this model. Additionally, depletion studies confirmed that these cells were indispensable to this treatments effect. Hence, our findings highlight the influential role of neutrophils to the treatment response of SBRT and IL-12 immunotherapy in PDAC.

