## Supplementary data for "Improving SBRT by re-wiring immunosuppressive neutrophils in murine pancreatic ductal adenocarcinoma"

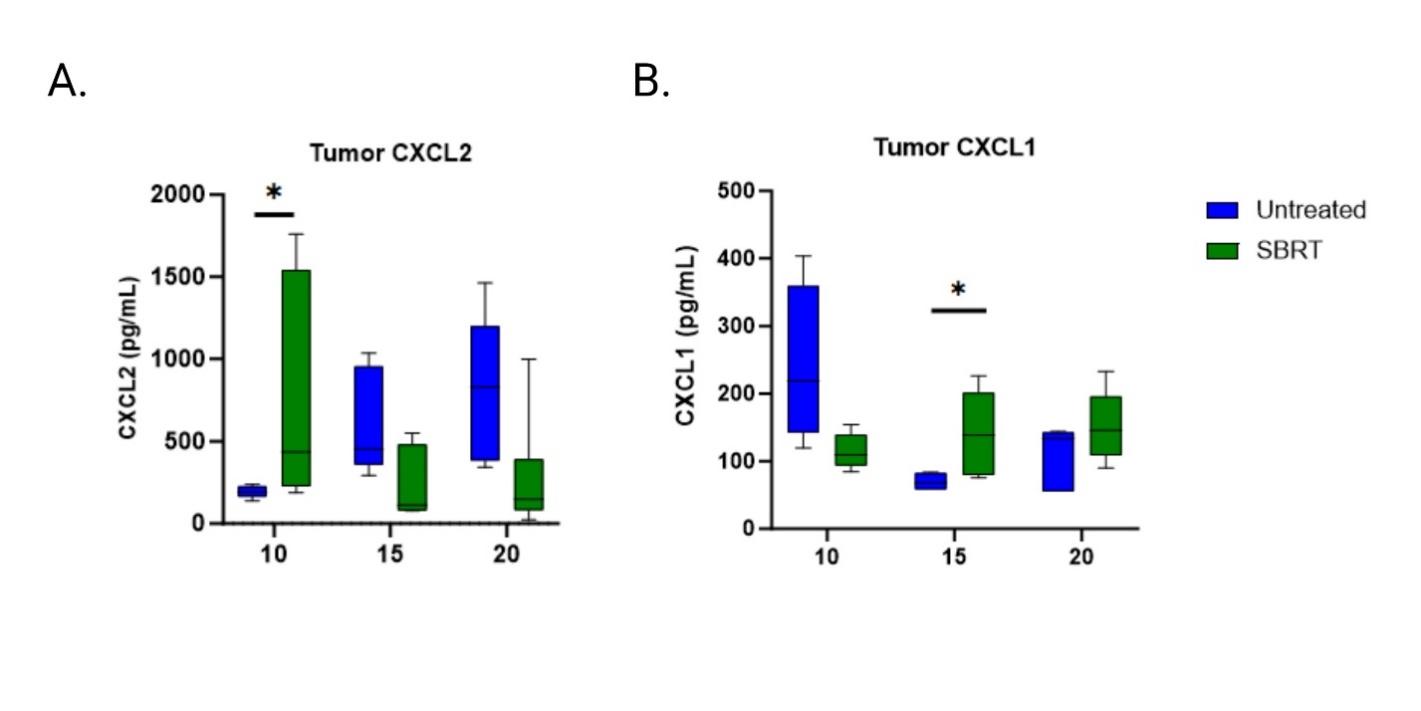
**Supplementary Figure 1. ELR+ chemokines are elevated in irradiated tumors with SBRT**

Box and Whisker plots illustrating intratumoral levels of **A**. CXCL2 and **B**. CXCL1 identified by Luminex assay (n=5). *P<0.05. Data analyzed by Mann-Whitney U tests.

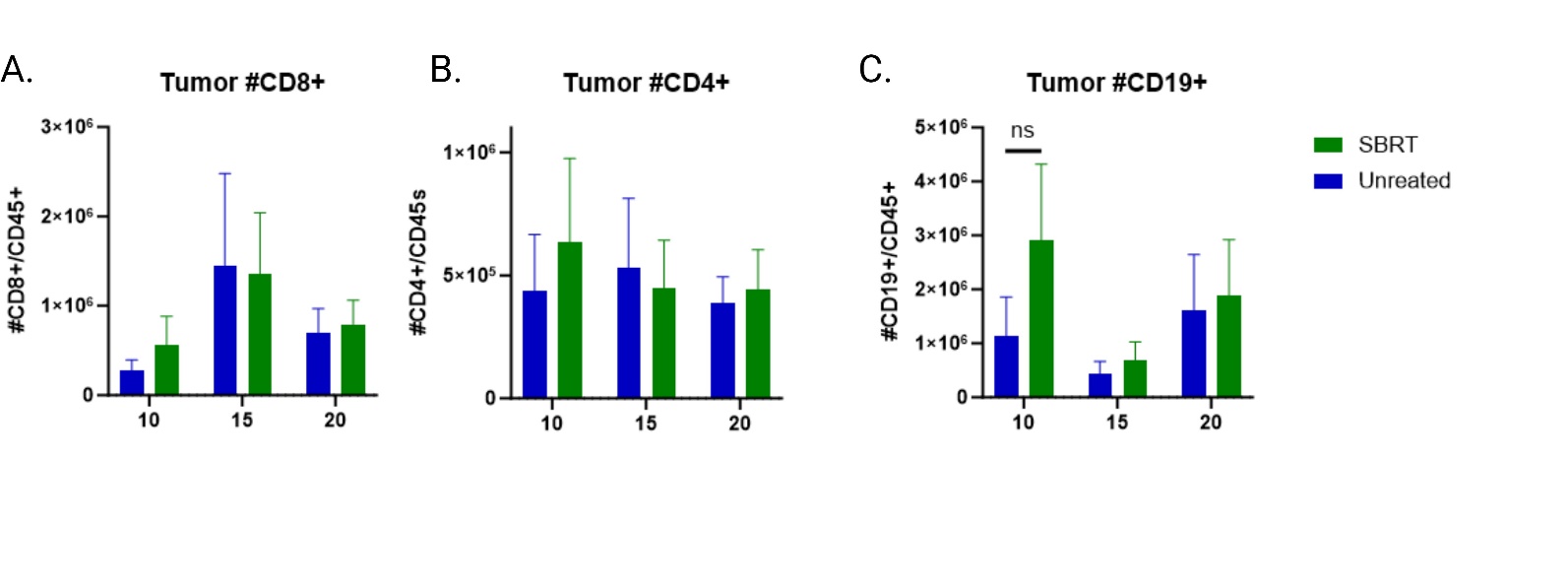
**Supplementary Figure 2. Lymphoid cell populations are unchanged following SBRT.**

**A**. CD8 T cell levels after SBRT. **B**. CD4 T cell levels after SBRT. **C**. CD19+ B cells after SBRT. Data shown as Mean + SD, *p<0.05. n=5. Data analyzed by non-parametric Mann-Whitney U test.
